# Spontaneous mentalizing after early interpersonal trauma: Evidence for hypoactivation of the temporoparietal junction

**DOI:** 10.1101/487363

**Authors:** Anna R. Hudson, Charlotte Van Hamme, Lien Maeyens, Marcel Brass, Sven C. Mueller

## Abstract

Experience of interpersonal trauma and violence alters self-other distinction and mentalizing abilities (also known as theory of mind, or ToM), yet little is known about their neural correlates. This fMRI study assessed temporoparietal junction (TPJ) activation, an area strongly implicated in interpersonal processing, during spontaneous mentalizing in 35 adult women with histories of childhood physical, sexual, and/or emotional abuse (childhood abuse; CA) and 31 women without such experiences (unaffected comparison; UC). Participants watched movies during which an agent formed true or false beliefs about the location of a ball, while participants always knew the true location of the ball. As hypothesized, right TPJ activation was greater for UC compared to CA for false versus true belief conditions. However, posttraumatic stress symptomatology (PTSS) appeared to play a role in driving the neural effect. In addition, CA showed increased functional connectivity relative to UC between the rTPJ and dorsomedial prefrontal cortex. Finally, the agent’s false belief about the presence of the ball speeded participants’ response (ToM index), but without group differences. These findings highlight that experiencing early interpersonal trauma can alter brain areas involved in the neural processing of ToM and perspective-taking during adulthood.

---

More than one in three women in a recent survey reported experiencing physical, sexual, and/or psychological violence in childhood [1]. Importantly, experiencing early traumatic life events like childhood abuse (CA) disrupts the normal developmental trajectory and alters mentalizing abilities, jeopardizing social interaction later in life; CA impairs cognitive perspective-taking (cf. [2]), reduces the ability to recognize and correctly interpret others’ emotions [3], and leads to less effective use of conflict-resolution strategies [4] in children. Nevertheless, while evidence mounts for the long-term consequences of CA on the neurobiology of affective and cognitive functioning [5-8], the long-term influence on mentalizing abilities is virtually unknown.

Despite the fundamental necessity of mentalizing abilities for daily life, neurobiological research examining the effects of interpersonal trauma on social cognition is scarce, and has mainly been conducted in psychiatric populations. To the best of our knowledge, only one imaging study has investigated the neural correlates of cognitive ToM in CA survivors: Quidé and colleagues [9] reported temporoparietal junction (TPJ) hypoactivation in a sample of previously maltreated adults with psychotic disorder. However, this study focused on the role of CA in impaired ToM in schizophrenia, and did not include healthy controls. By comparison, a study in adolescents [10] documented a link between abuse severity and inferior frontal gyrus (IFG) activation but focused on an affective ToM task, namely the Reading the Mind in the Eyes Task (RMET). Abu-Akel and Shamay-Tsoory [11] have proposed a model dissociating cognitive and affective ToM, concluding that there is evidence that these are separate processes with distinct yet connected neural networks. Meta-analytic findings support this model suggesting that cognitive ToM tasks elicit more bilateral TPJ and precuneus activation whereas affective tasks involve the bilateral IFG and posterior medial frontal cortex [12]. Furthermore, both of the above studies employed explicit – rather than spontaneous – mentalizing tasks; i.e., while they may be informative regarding *ability* to mentalize in maltreated populations, evidence is lacking regarding *propensity* to mentalize (see [13] for a theoretical article regarding *ability* versus *propensity*). In addition, because adults can easily solve explicit ToM tasks, recent neuroimaging work has developed implicit ToM tasks [14], in which participants are not asked explicitly to take the perspective of another into consideration. These tasks have been further validated in clinical settings with populations with ToM deficits such as autism [15]. Together with earlier work [16-19], these studies [14,15] highlight a particularly crucial role of the right TPJ as the main region of interest for these kind of tasks.

Therefore, the present study sought to identify the impact of CA on spontaneous cognitive ToM and rTPJ functioning during adulthood. Based on behavioral [20,21] and neural [9] evidence, this study tested the direct hypothesis that exposure to CA will lead to hypoactivation of the rTPJ, indicative of compromised ToM, in a community sample of adult CA survivors. To this end, women with experience of CA and women without such histories completed a well-validated implicit ToM task [14,15,22,23] during fMRI.

## Methods

### Participants

Thirty-five adult women with a history of childhood abuse (CA; mean age = 36.79 years, *SD* = 12.04) and 40 adult women without such history (unaffected comparison, UC; mean age = 35.64 years, *SD* = 11.50) participated in exchange for €30 (Table 1). Inclusion criteria for CA were experience(s) of physical, sexual, and/or emotional abuse occurring before age 17 (Supplementary Table S1.1). Inclusion criteria for UC were no experience of childhood trauma and no experience of interpersonal trauma (e.g. emotional abuse, physical/sexual assault, etc.) later in life. Because it emerged during questionnaires that nine women initially recruited as UCs had had such experiences, these participants were excluded, resulting in a final dataset of 31 adult women with no history of childhood or interpersonal trauma (mean age = 36.51 years, *SD* = 11.46). Rerunning analyses with these women included did not substantially alter findings. Inclusion criteria for both groups were: MRI compatibility (i.e., no pregnancy or metal implants), fluency in Dutch, no history of severe head trauma or severe neurological condition, normal or corrected-to-normal vision, and being 18-60 years old. Participants from both groups were recruited without regard to psychiatric history, as childhood maltreatment is a risk factor for a wide range of psychological disorders [24] and including only healthy survivors of childhood abuse could bias results [25](Table1, Supplementary material).

Participants were recruited via self-help groups (CA only), flyers, social media, and the student pool of Ghent University and were matched for age, sex, handedness, and level of education (𝒳^2^_(2)_ = 1.96, *p* = .38). CA had significantly higher levels of self-reported empathy (*t*_(64)_ = 3.03, *p* = .004, *d* = 0.75), depression (*t*_(49.21)_ = 5.07, *p* < .001, *d* = 1.23), dissociation (*t*_(51)_ = 3.52, *p* = .001, *d* = 0.98), trait anxiety (*t*_(63)_ = 5.80, *p* < .001, *d* = 1.44), state anxiety (*t*_(64)_ = 4.61, *p* < .001, *d* = 1.14), and current psychopathology (*t*_(64)_ = 5.85, *p* < .001, *d* = 1.44) (Table 1). UC participants self-reported significantly higher levels of resilience (*t*_(64)_ = 3.16, *p* = .002, *d* = 0.78) but were equally likely to be taking psychotropic medication (𝒳^2^_(1)_ = 2.56, *p* = .11). The study was approved by the IRB of Ghent University Hospital and all participants provided written informed consent prior to commencing the study.

**Table 1.**
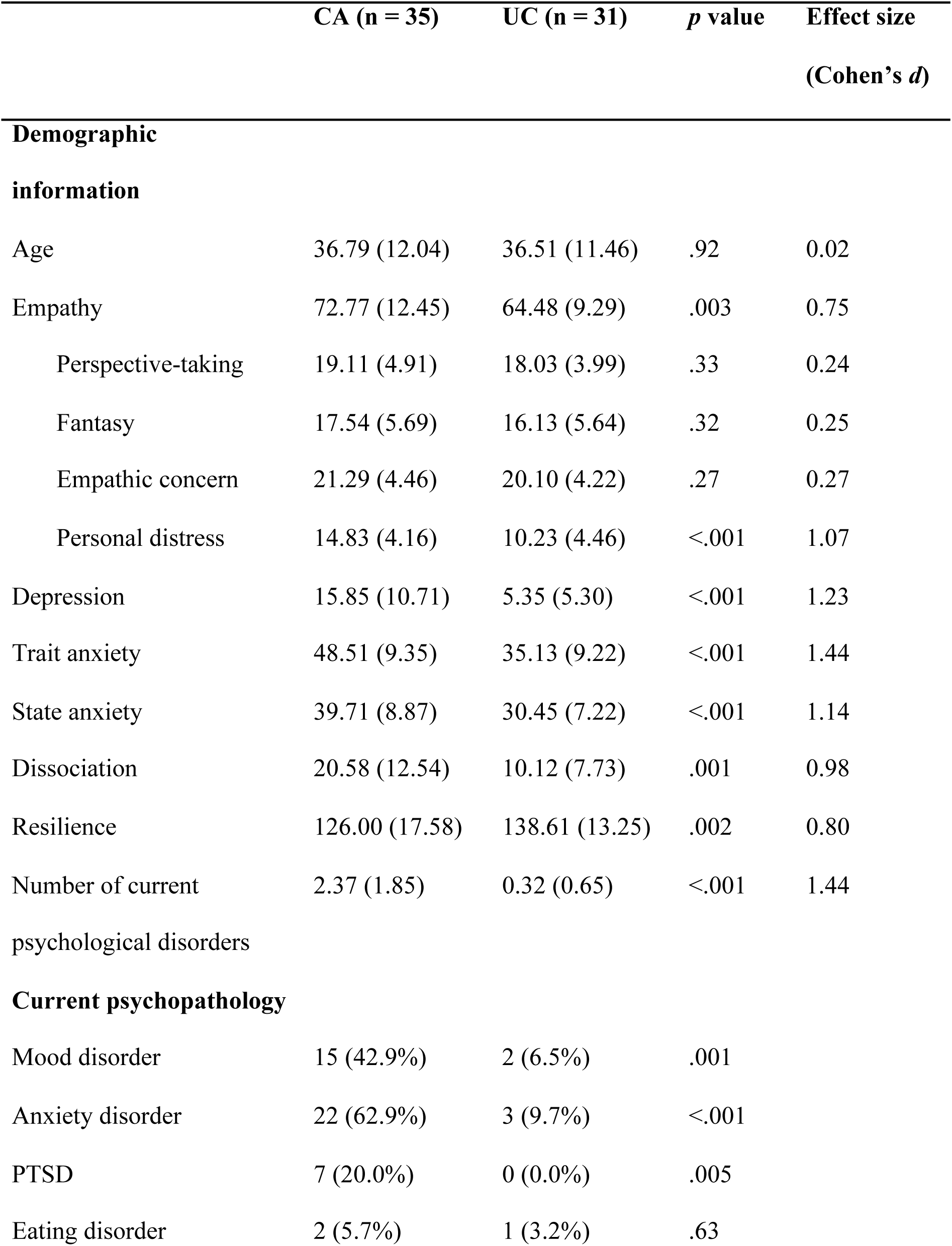
Sample demographics, current psychopathology (MINI), and mean scores on measures of empathy (IRI), depression (BDI), trait and state anxiety (STAI), dissociation (DES), and resilience (RS).

## Materials

### Implicit ToM task

An adapted version [26] of a previously validated task [22,23] was used to measure implicit (spontaneous) ToM. The task comprises two phases – a *belief phase*, where the beliefs of the participant and an onscreen agent (Buzz Lightyear; Toy Story, 1995) are manipulated, and an *outcome phase*, where participants are required to react to the presence of a ball. The task consists of eight different movies and was presented using Presentation 18.2 (NeuroBehavioral Systems, Inc.).

At the beginning of each movie, Buzz Lightyear enters the scene and rolls a ball on a table with an occluder. Four different scenarios are then possible (Figure 1). In two of the scenarios, the ball stops behind the occluder. In the other two scenarios, the ball comes to rest off-screen. The agent then leaves the scene. Subsequently, in half of the scenarios, the ball rolls again and changes location (i.e., if it were originally behind the occluder, it rolls and comes to a final stop off-screen and vice versa). In the other two scenarios, the ball remains stationary and does not change location. The agent then returns to the scene. Thus, while the participant always knows the ball’s true location, in half of scenarios, the agent falsely believes the ball to be somewhere else to where it really is. At the end of each movie, the occluder falls. Each of the four scenarios was created with a “ball present” and “ball absent” ending, resulting in eight different movies.

Participants’ task was to press a button as quickly and as accurately as possible if the ball was present after the occluder fell. In reality, 50% of movies ended with the ball present, regardless of participant’s or agent’s belief. Participants were also required to press another button when the agent left the scene, to ensure that they paid attention to the whole movie. Crucially, participants were given no instructions to pay attention to the agent’s beliefs.

**Figure 1.**
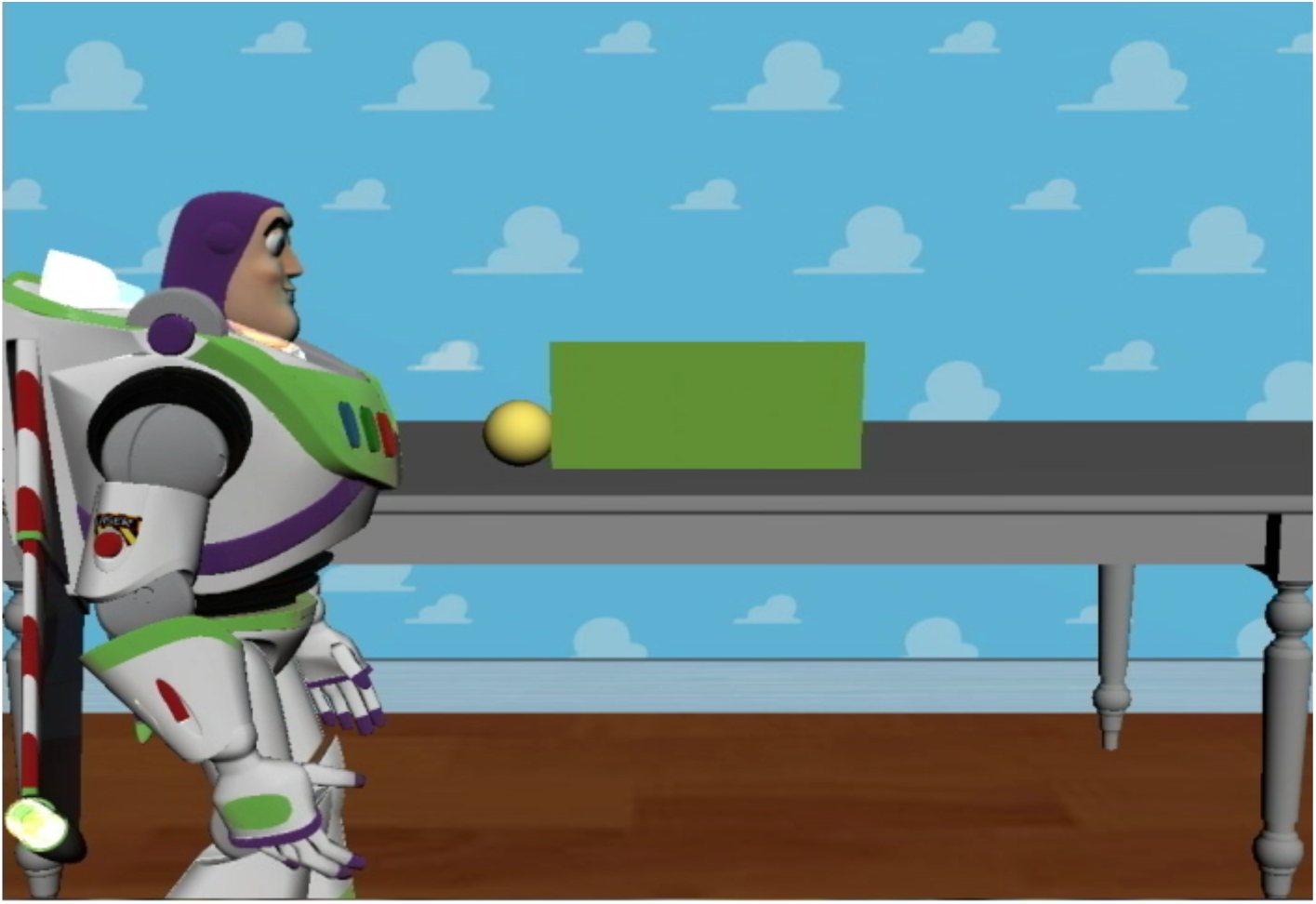
Screenshot of a sample trial (for description see text).

### Theory of Mind localizer

To identify suitable mentalizing-related brain regions and cohort-specific coordinates for these, including the rTPJ (primary ROI analysis), and to maintain independence of analysis, a Dutch translation of a well-validated ToM localizer was used [27]. Participants read 20 short stories pertaining either to: characters with false beliefs (“false belief” condition), or to inanimate objects such as maps or photographs which display false information (“false photograph” condition). Following each story, participants read a statement on that story. Half of the stories (10) belonged to the “false belief” condition whereas the other half (10) belonged to the “false photograph” condition. Participants then read a statement on that story and responded using an MRI compatible response box (Cedrus)(index finger = true statement, middle finger = false statement).

### Questionnaires

#### SLESQ

The 11-item Stressful Life Events Screening Questionnaire (SLESQ; [28]) was used to assess previous traumatic exposure. Sub-items ask for additional details such as age at the time of the experience, frequency, and duration of the trauma. Participants were classified as MW if they positively answered items 5 (rape), 6 (sexual assault), 7 (childhood physical abuse), and/or 9 (emotional abuse), and if their age at onset was below 17 years. This cut-off was chosen to keep similarity with other questionnaires assessing childhood abuse, such as the Childhood Trauma Questionnaire [29]. Cronbach’s alpha (α) for the present sample was .76, indicating good internal consistency.

#### IRI

The 28-item Interpersonal Reactivity Index [30] (IRI; Dutch translation [31]) measures empathic responsiveness (α = .79). It assesses four aspects of empathy: two cognitive (perspective-taking and fantasy) and two affective (empathic concern and personal distress).

#### BDI-II

The 21-item Beck Depression Inventory-II [32] (BDI-II; Dutch translation [33]) measures depressive symptoms covering cognitive, affective, and somatic aspects of depression (α = .94).

#### STAI (trait and state)

The State-Trait Anxiety Inventory [34] (STAI; Dutch translation [35]) measures both state anxiety (20 items, α = .92) and trait anxiety (20 items, α = .94).

#### DES

The 28-item Dissociative Experiences Scale [36] (DES; Dutch translation [37]) measures the extent to which respondents experience dissociative symptoms such as depersonalization, derealization, and disturbances in memory and identity in their daily life (α = .90).

#### RS

The 25-item Resilience Scale [38] (RS) measures mental resilience and adaptability (α = .86).

### Measure of psychopathology

#### MINI

The Mini International Neuropsychiatric Interview [39] (MINI; Dutch version [40]) was administered by trained clinical psychology masters students. It assesses current and lifetime histories of Axis I disorders plus one Axis II disorder, antisocial personality disorder, based on DSM-IV criteria. Due to much previous research on the link between experience of CA and later borderline personality disorder (BPD) [41,42], it was decided to include an additional MINI-style subsection assessing BPD symptomatology. Items were designed based on DSM-IV criteria, translated into Dutch, and back-translated. It was decided not to use a more thorough diagnostic interview (such as the Structured Clinical Interview for DSM-5 (SCID-5) [43] or the Clinician Administered PTSD Scale (CAPS-5) [44]) due to time constraints; furthermore, as our main focus was the influence of childhood trauma rather than psychopathology, the MINI was deemed sufficient.

## Procedure

During fMRI participants completed the implicit ToM task (∼20 mins duration with a short break in the middle) followed by the ToM localizer (∼10 mins duration) in fixed order. For the implicit ToM task, participants were instructed to respond with their left index finger as quickly and as accurately as possible when the agent left the scene and to respond with their right index finger as quickly and as accurately as possible if the ball was present at the end of the movie (when the occluder fell). Prior to the task proper, feedback was given on four practice trials. The task consisted of eight movies of each condition, resulting in 64 experimental trials, split into two blocks, and presented in randomized order. Each trial lasted 13,800 ms. In each trial, the occluder fell at 13,250 ms. Between trials, there was a jitter of variable duration during which a black screen was displayed.

For the ToM localizer, each trial comprised a short story displayed for 10 s, followed by a statement which was also displayed for 10 s. Beneath the statement, the words “true” and “false” (Dutch: “juist” and “onjuist”) were displayed on either side of the screen to remind participants which finger (left and right index finger, respectively) to respond with. The task consisted of 20 trials (10 per condition), presented in fixed order. Between trials, there was a jitter of variable duration during which a white fixation cross was displayed on a black background.

Following scanning, participants completed the questionnaires outside the MRI scanner. Lastly, the MINI was conducted. Participants were then paid, debriefed, and thanked.

### fMRI data acquisition

Images were acquired using a 3T Magnetom Siemens TrioTim MRI scanner at Ghent University Hospital. First, a T1-weighted high resolution anatomical scan was performed (repetition time [TR] = 2250 ms, echo time [TE] = 4.18 ms, image matrix = 256 × 256, field of view [FOV] = 256 mm, flip angle = 9°, slice thickness = 1 mm, voxel size = 1.0 × 1.0 × 1.0 mm, number of slices = 176). Functional images were acquired using a T2*-weighted Echo Planar Images (EPI) sequence (TR = 2000 ms, TE = 28 ms, image matrix = 384 × 384, FOV = 224 mm, flip angle = 80°, slice thickness = 3.0 mm, voxel size = 3.5 × 3.5 × 3.0 mm, number of slices = 34).

### fMRI data processing

Data were preprocessed with SPM12 (Wellcome Department of Cognitive Neurology, London, UK) implemented in MATLAB R2012b (Mathworks, Inc.). The first five volumes of all EPI sequences were discarded due to scanner calibration. Data preprocessing began with coregistering functional images with the corresponding participant’s structural image. Next, coregistered images were realigned and slice time correction applied, followed by again coregistering the images with their respective structural image. Structural images were then segmented, and the parameters generated during segmentation used to normalize the images to standard MNI space, after which voxel size was 2 × 2 × 2 mm. Finally, images were spatially smoothed with a Gaussian kernel of 8 mm (full-width at half maximum). The high-pass filter was set at 128 Hz.

Motion correction was applied following preprocessing using Bramila Framewise Displacement (Brain and Mind Lab, Helsinki, Finland). No participant had a mean framewise displacement above 0.5 mm (recommended for functional connectivity analyses [45,46]), with overall mean displacement 0.09 mm for CA (*SD* = 0.05) and 0.08 mm for UC participants (*SD* = 0.03).

### fMRI data analysis

All trials were modeled in SPM regardless of whether participants responded correctly. First-level statistical analyses were performed using the general linear model (GLM).

#### ToM localizer

For the ToM localizer, story and statement phases were conflated and analyzed together. First-level models contained separate regressors for each condition (“false belief” and “false photograph”) as well as subject movement parameters. The second-level model contained three regressors of no interest, namely depression score (BDI-II), trait anxiety score (STAI-T), and age. In order to identify primary (rTPJ) and secondary ToM regions, a one-sample *t*-test using the whole group was performed on the false belief > false photograph contrast at the whole brain level (Supplementary Table S2). Coordinates of peak activation for these regions (corrected for multiple comparisons (FWE) and thresholded at *p* < .05 and cluster size *k* > 10) were then used in the ROI analyses for the implicit ToM task.

#### Implicit ToM task (belief phase)

For the implicit ToM task, only the belief phase was analyzed (thus conflating “ball present” and “ball absent” conditions into one, and resulting in four conditions). First-level models contained separate regressors for each condition as well as six movement parameters. In order to identify regions involved in ToM, a false belief (FB) > true belief (TB) contrast was generated. In this contrast, conditions where the agent falsely believed the ball to be in a different location to its true location (i.e., where the agent’s belief differed from the participant’s) were classified as “false belief” and conditions where the participant and the agent both believed the ball to be in the same location were classified as “true belief”. In order to investigate differences in spontaneous ToM between CA and UC, first-level contrasts were entered into a second-level model and a UC > CA contrast was created. Regressors of no interest were: depression score (BDI-II), anxiety score (STAI-trait), and age. A primary a priori-defined ROI analysis was carried out on the rTPJ and other regions (secondary analysis) found to have been activated by the ToM localizer. These regions are also commonly reported in meta-analyses to be involved in mentalizing [12,47,48]. In order to ensure independence of analysis, coordinates for ROI analyses (6 mm spheres) were taken from coordinates of peak activation in the ToM localizer. As the temporal poles are well-defined anatomically, here we instead used masks created with WFU PickAtlas [49,50]. Data were analyzed using the SPM toolbox MARSBAR [51]. Beta values for regions of interest were extracted and correlated with participants’ *ToM index* (see behavioral data analysis) and questionnaire data. As a manipulation check (post-hoc), we examined whether regions elicited during the ToM task overlapped with our ROIs from the ToM localizer, which they did (Supplementary Table S3), suggesting that the localizer was valid in eliciting cognitive ToM regions.

In order to investigate functional connectivity between ToM regions, VOI information were extracted from the rTPJ and FB > TB contrasts were generated. It was decided to limit analyses to this region, as this was the main focus of this research and furthermore the most consistently activated in cognitive ToM tasks [12]. Psychophysiological interaction (PPI) analyses were then performed on this contrast, again using depression score, trait anxiety score, and age as regressors of no interest. Furthermore, in order to investigate connectivity specifically between ToM regions, a priori-defined ROI analyses were performed on the same regions as in the functional analyses (6 mm spheres). Data were analyzed using the SPM gPPI and MARSBAR toolboxes [51,52].

### Behavioral data analysis (outcome phase)

As in previous work [14,15,23,26], participants’ *ToM index* on the implicit ToM task was computed by calculating the difference in RT during the outcome phase between the P-A-condition (neither participant nor agent expect to see the ball) and P-A+ condition (where the participant knows the ball should not be present but also knows that the agent believes the ball will be). In this way, it is possible to see how the agent’s perspective can spontaneously influence RT. A positive *ToM index* would thus indicate a facilitating effect. Participants were excluded from analyses if their *ToM index* was more than 2.5 standard deviations from the group mean (n_MW_ = 1; n_UW_ = 0). Independent *t*-tests were used to compare group performance, with Cohen’s *d* as a measure of effect size. Furthermore, questionnaire data were correlated with the *ToM index* (Bonferroni-corrected). All analyses were performed in IBM SPSS Statistics 25.0 with *p* < .05 (two-tailed). Behavioral data for the ToM localizer were not analyzed as the purpose of this task was to generate coordinates for fMRI ROI analyses.

## Results

### Regions of interest

First, regions involved in ToM including the rTPJ were identified using the ToM localizer. Analysis revealed significant clusters located in the bilateral TPJ, precuneus, dmPFC, and bilateral MTG/temporal poles (Supplementary Table S2), with no significant group differences.

### Neural activation during belief phase

#### Primary ROI analysis: rTPJ

As expected, analysis of the rTPJ (peak MNI xyz: 46 - 58 20) revealed that UC (*n* = 31) had significantly more activation in this ROI than CA (*n* = during the belief phase of the ToM task (*t*_(58)_ = 1.70, *p* = .048) with a medium effect (*d* = 0.45) (Figure 2).

**Figure 2.**
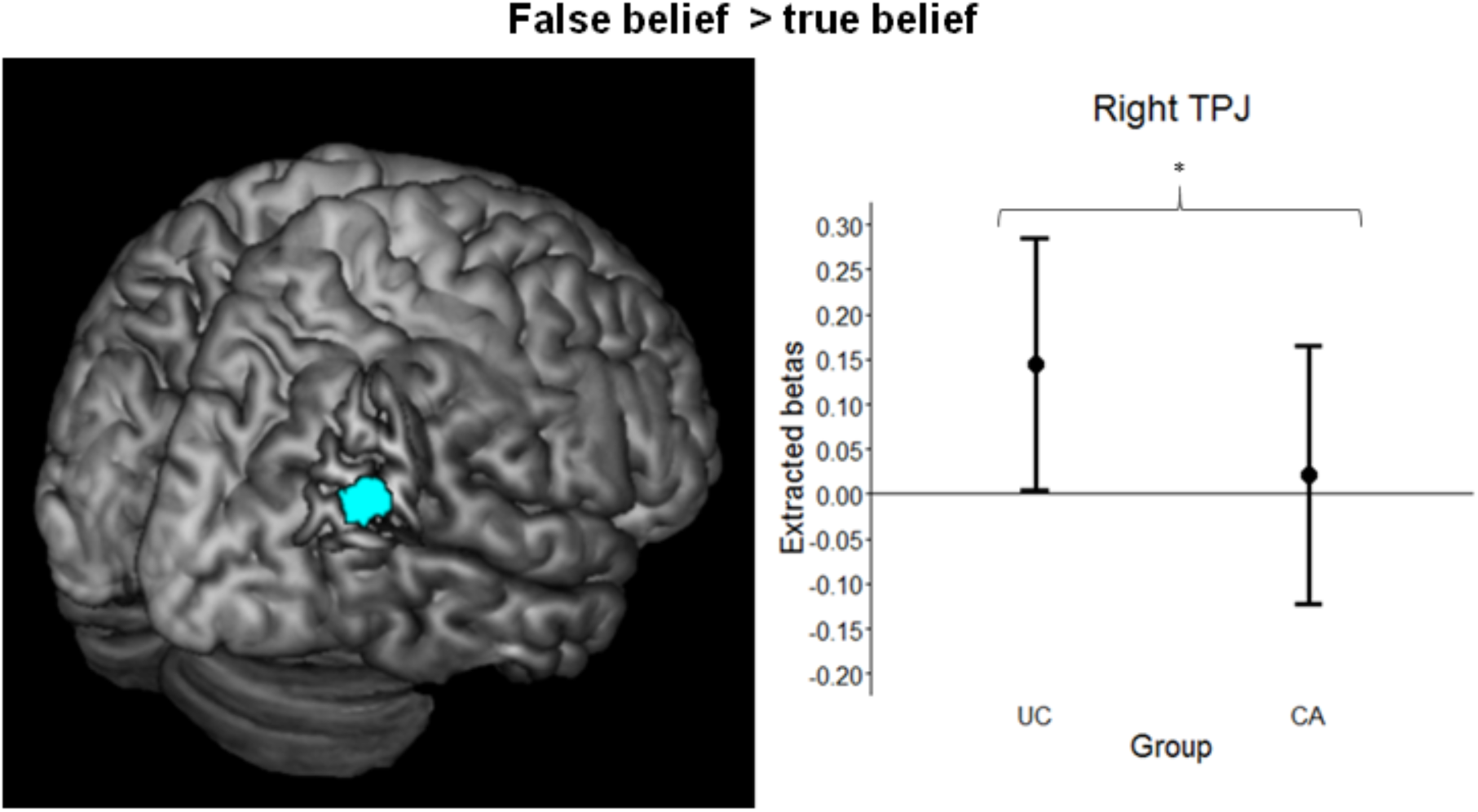
Right temporoparietal junction (TPJ) region of interest (6 mm sphere) on a standard brain (left panel). Mean extracted betas and 95% confidence intervals for the rTPJ in women with childhood abuse history (CA, *n* = 35) and unaffected comparison women (UC, *n* = 31). False belief (FB) > true belief (TB) (right panel). * = *p* < .05.

#### Secondary ROI analysis

ROI analyses on other regions generated by the localizer (Figure 3) revealed a similar, but reduced, effect in the lTPJ (peak MNI xyz: -48 -56 24) (*t*_(58)_ = 1.12, *p* = .13, *d* = 0.29). RTPJ activation was significantly positively correlated with lTPJ activation (CA: *r*_(35)_ = .53, *p* = .001; UC: *r*_(31)_ = .48, *p* = .006) but not with questionnaire data or self-reported empathy. Analysis of the dmPFC (MNI xyz: -10 54 28), left MTG (MNI xyz: -56 -2 -18), and left temporal pole (Brodmann area 38) revealed significantly higher activation in UC compared to CA, all with medium effect sizes (dmPFC: *t*_(58)_ = 2.03, *p* = .02, *d* = 0.53; MTG: *t*_(58)_ = 1.92, *p* = .03, *d* = 0.50; temporal pole: *t*_(58)_ = 1.62, *p* = .06, *d* = 0.43). Analysis of the right MTG, right temporal pole, and precuneus ROIs revealed no significant group differences.

**Figure 3.**
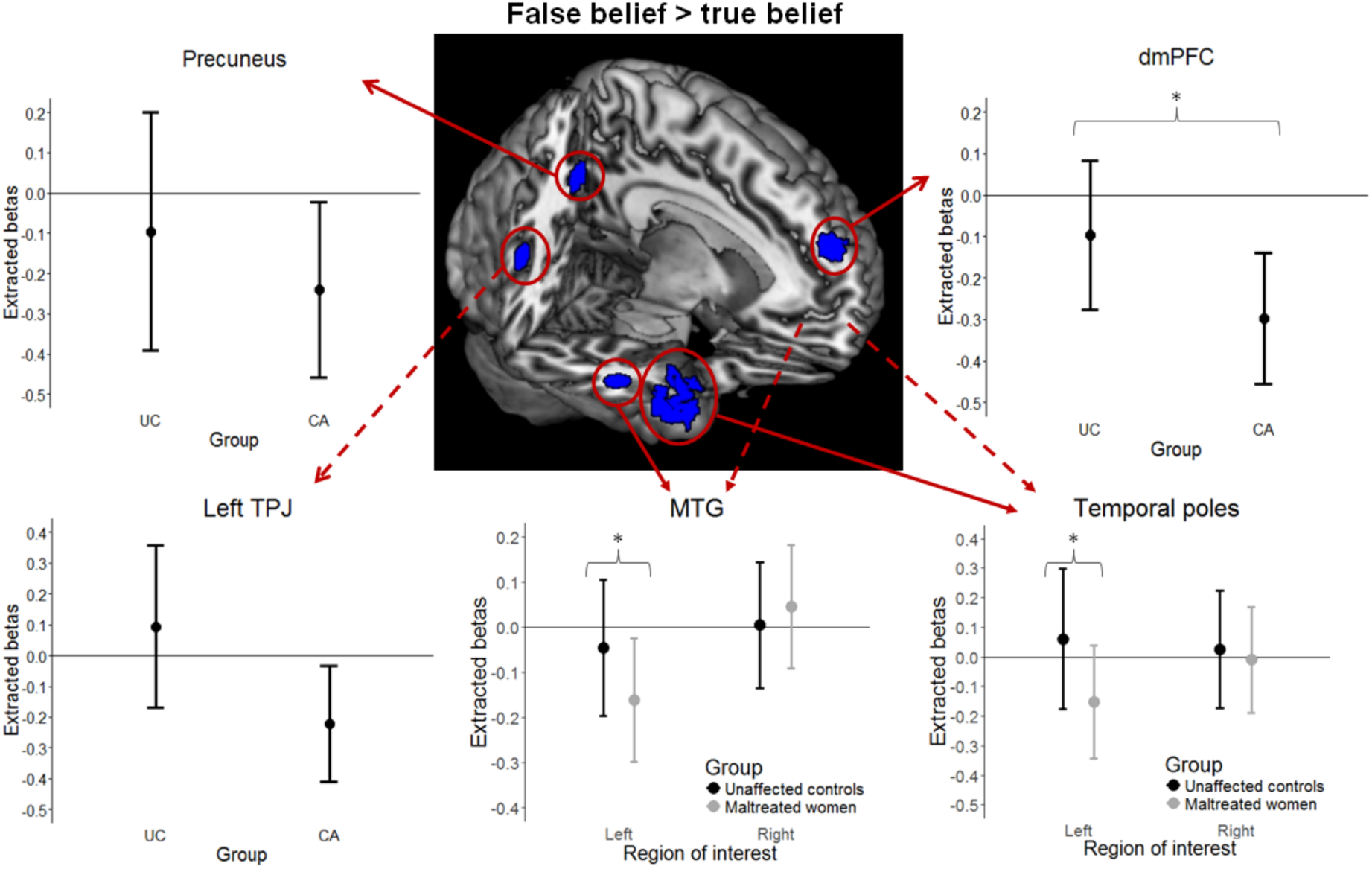
Secondary regions of interest on a standard brain. ROIs were 6 mm spheres except for the temporal poles, which were defined anatomically as Brodmann area 38. Mean extracted betas and 95% confidence intervals for secondary ROIs in women with childhood abuse history (CA, *n* = 35) and unaffected comparison women (UC, *n* = 31). Dashed lines indicate bilateral ROI. The right TPJ is displayed. *Key:* TPJ = temporoparietal junction; dmPFC = dorsomedial prefrontal cortex; MTG = middle temporal gyrus. * = *p* < .05.

### Comorbid psychopathology

#### PTSD

To investigate the influence of PTSD symptomatology (PTSS), we divided CA into non-PTSD and (sub-)clinical PTSD (as per the MINI) and reran models comparing UC > CA-non-PTSD and UC > CA-PTSD. CA were categorized as “non-PTSD” if they endorsed only one of the two MINI PTSD screening questions (*n* = 22). CA were categorized as “subclinical PTSD” if they endorsed both screening questions but did not reach clinical threshold on the follow-up questions. As there were few CA with (sub-)clinical levels of PTSS (n_clinical_ = 7; n_subclinical_ = 6), we combined these two groups; furthermore, visual inspection revealed little difference between CA-PTSD and CA-subclinical PTSD. Findings between UC and CA-non PTSD were no longer significant for the lTPJ or rTPJ (*p*s > .05), yet remained significant between UC and CA-PTSD for the rTPJ (*t*_(36)_ = 1.68, *p* = .05, *d* = 0.56), and to a lesser extent for the lTPJ (*t*_(36)_ = 1.16, *p* = .13, *d* = 0.38) (Figure 4).

#### BPD

To investigate the influence of BPD, we reran models comparing UC > CA-non-BPD (*n* = 23) and UC > CA-BPD (*n* = 12). Findings between UC and CA-non-BPD held for the l/rTPJ (rTPJ: *t*_(35)_ = 1.95, *p* = .03, *d* = 0.66; lTPJ: *t*_(35)_ = 1.59, *p* = .06, *d* = 0.54), but disappeared for UC > CA-BPD (*p*s > .05).

#### Other psychopathology

With regards to other psychopathology, there was no influence of depression and anxiety, alcohol and substance disorders, or psychosis. For an in-depth description of analyses, see Supplementary 3.1.

**Figure 4.**
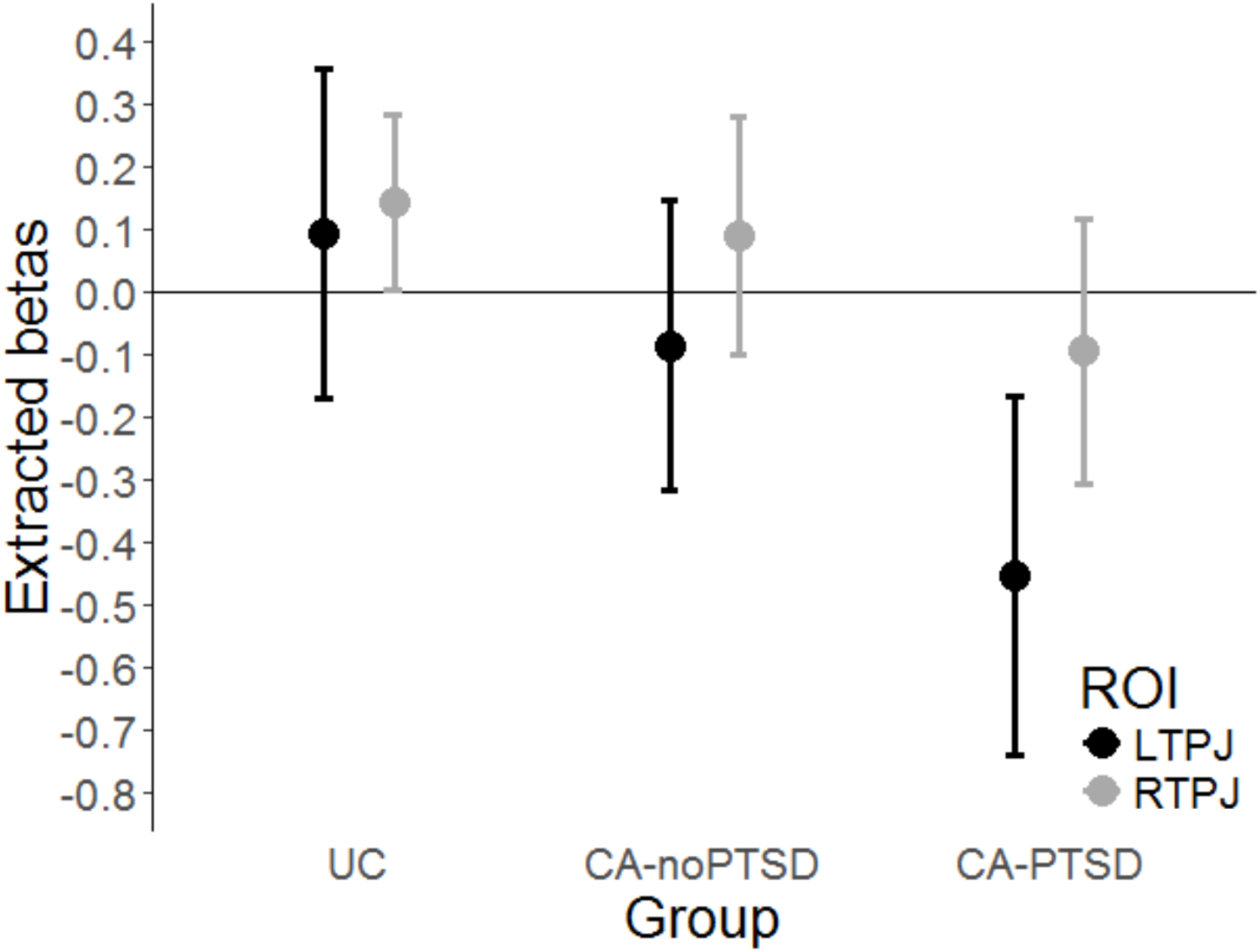
Mean activation of the left and right temporoparietal junction and 95% confidence intervals for PTSD diagnoses. *Key:* UC = unaffected comparison women (*n* = 31); CA-no PTSD = women with childhood abuse history who endorsed only one of the two PTSD screening items and thus were not asked the follow-up questions (*n* = 22); CA-PTSD = women with childhood abuse history who reached a (sub-)clinical diagnosis of PTSD, i.e. who endorsed both PTSD screening items and were asked the follow-up questions (*n* = 13). CA-subclinical PTSD and CA-clinical PTSD categories were collapsed due to minimal difference in activation. * = *p* < .05.

### Functional connectivity during belief phase

#### Psychophysiological interactions

CA showed greater functional connectivity (FC) between the rTPJ and dmPFC ROI during the ToM contrast (FB > TB) relative to UC (*t*_(58)_ = 2.43, *p* = .009, *d* = 0.64). No other effects were significant. Follow-up one sample *t*-tests assessing directionality revealed that CA had significant positive FC between the rTPJ and dmPFC (*t*_(58)_ = 1.92, *p* = .03, *d* = 0.50) whereas UC had trending negative FC (*t*_(58)_ = 1.53, *p* = .07, *d* = 0.40).

#### Influence of psychopathology

Due to the influence of PTSS and BPD in the functional analyses, we also examined the impact of these on FC between the rTPJ and dmPFC. Diverging from the functional results, CA-non-PTSD (*n* = 22) showed considerably higher FC in comparison to UC (*t*_(45)_ = 3.39, *p* < .001) with a large effect size (*d* = 1.01), with no difference between UC and CA-PTSD (*n* = 13). Conversely, both CA with (*n* = 12) and without BPD (*n* = 23) showed greater FC with the dmPFC compared to UC (CA_BPD_: *t*_(35)_ = 2.30, *p* = .01, *d* = 0.78; CA_non-BPD_: *t*_(46)_ = 2.01, *p* = .02, *d* = 0.59).

#### Behavioral results during outcome phase

There was no significant difference in *ToM index* between UC and CA (*t*_(62)_ = 0.98, *p* = .33, *d* = 0.25) (Supplementary Table S4) and no correlations between the *ToM index* and the questionnaire data.

## Discussion

This study tested - and confirmed - the simple hypothesis that experience of early life maltreatment is associated with reduced activation of the TPJ, a core structure in social cognition and ToM. Consistent with the central hypothesis, women with CA experiences showed less rTPJ activation during the cognitive mentalizing task comparative to women without such experiences. Against expectations, no significant behavioral differences emerged. However, task differences appeared to be driven by comorbid PTSS.

In keeping with our hypothesis, previous studies on childhood maltreatment in psychiatric populations [9], and populations known to have mentalizing deficits (e.g. autism spectrum disorder [15]; schizophrenia [53]), CA displayed rTPJ hypoactivation while observing an agent form false beliefs during a mentalizing task. Furthermore, such hypoactivation was also observed in other crucial regions associated with ToM, namely the dmPFC, the left MTG, and the left temporal pole. The present data support the hypothesis that women who have been maltreated during childhood show aberrant functioning of the ToM network, specifically while computing other people’s beliefs when these differ from their own. However, PTSS caused by this maltreatment experience appeared to drive the effect.

Indeed, exploratory analyses revealed that the more severe the PTSS (as per the MINI), the lower TPJ activation was, with CA classified as having (sub-)clinical PTSD faring worse than CA with little to no PTSS. Thus, TPJ hypoactivation may be related to the degree to which the individual was psychologically affected by their traumatic experience. Our findings thus align with a previous behavioral study by Nazarov and colleagues [21], who found adult women with CA-related PTSD to have reduced ToM relative to unaffected women. The present results build upon their findings and show that women with CA-related PTSS also have reduced mentalizing abilities in comparison to women who experienced CA but did not develop (sub-)clinical PTSD. To our knowledge, no other studies have assessed mentalizing deficits in PTSD sufferers compared to trauma-exposed controls, neither behaviorally nor neurologically. Thus, while the present data are preliminary, they raise interesting questions regarding how trauma-exposed individuals without PTSD differ both from non-trauma-exposed individuals and PTSD sufferers.

Yet, a notable discrepancy emerged when examining functional connectivity. Here, CA without PTSS showed larger FC relative to comparisons, while CA-PTSD did not show such an effect. This may suggest an interesting distinction between women who develop PTSS and those who do not. Arguably, and speculatively at present, CA without PTSS might be able to utilize compensatory mechanisms that can maintain adequate levels of ToM processing within the network. By contrast, women with PTSS, who were more likely to display hypoactivation of the TPJ and no such increased FC, may no longer be able to instil such compensatory mechanisms, potentially because of fundamental changes to the ToM network. On the neural level, it seems that CA with and without PTSS can be differentiated in their ToM processing. Further study will need to identify the behavioral consequences of this effect.

Interestingly, presence of BPD appeared to have an opposite effect. While CA without BPD showed hypoactivation relative to UC, notably there emerged no such differences between CA with BPD and UC. Previous studies have found BPD sufferers relative to healthy controls to over-interpret/attribute mental states (“overmentalize”) on ToM tasks [54-57] and show hyperactivation in key mentalizing brain regions, including the TPJ [55,58].

Furthermore, in recent behavioral [59] and neural [60] studies, Abu-Akel and colleagues posited that when an individual has comorbid autism (associated with undermentalizing) and psychotic symptomatology (related to overmentalizing, similarly to BPD), it may be possible that a “normalizing effect” occurs. This could therefore suggest that the diametrically opposed influences of CA and BPD resulted in such a “normalizing effect” in our own sample. However, this has to be taken with great caution.

Regardless, this study has important clinical implications. It appears that women with a history of CA, and specifically those with PTSS, are uniquely at risk of altered perspective-taking abilities well into adulthood. This would therefore indicate that individuals seeking psychotherapy who have experienced CA will require a specialized approach. This is of particular importance, as a recent study suggests that assaulted adolescent girls respond differently to trust violations, possibly due to alterations in their mentalizing abilities [61]. Tailored therapy appears warranted to help improve such abilities, in order to reduce risk of revictimization. Furthermore, reduced social competence has been linked to poorer social support networks in adolescents [62] and adults [63,64], which in turn is associated with a higher risk for psychopathology in maltreated women [65-67]. Moreover, our study highlights the impact PTSD can have on social cognitive skills, an impact which has the potential to create a vicious downward spiral and a topic remarkably under-researched.

Despite these important findings, there are some notable limitations. Firstly, our findings regarding PTSD must be interpreted with some caution as we had only a small number of CA reaching a (sub-)clinical diagnosis. Furthermore, we have limited information regarding symptom severity, as we assessed current psychopathology using a general psychopathology screening interview rather than a more specialized PTSD interview such as the CAPS [44]. However, the aim of this study was not to investigate the impact PTSS had on mentalizing and mentalizing-related brain regions, but rather to investigate the impact of experience of CA, which it did so successfully. Moreover, this study also gave valuable insight into a more representative sample of CA survivors – neither one extreme end of the spectrum (i.e., those with a clinical diagnosis) nor the other (i.e., those without any psychopathology). Secondly, our sample size consisted solely of female survivors of CA, so generalization to male survivors must be done cautiously. However, this was done for two reasons – firstly, there is evidence of neurological developmental differences between males and females [68] which could confound results in an unbalanced mixed-sex sample; and secondly, there is evidence of men and women responding differently, including on a neural level, to experience of abuse [69].

In conclusion, here we demonstrate hypoactivation of several key mentalizing brain regions in women with a history of childhood abuse. Clinically, this study emphasizes the importance of assessing individuals seeking psychotherapy for history of childhood abuse, as this may present a unique profile and set of risks independent of psychological diagnosis.

## Acknowledgements

The authors would like to warmly thank all women who participated in our research, as well as all organizations and groups who helped with recruitment. This work was supported by a 2-4 year grant (01J05415) from the Special Research Fund (BOF) at Ghent University to SCM and MB.

